# Inhibiting corticospinal excitability by entraining ongoing mu-alpha rhythm in motor cortex

**DOI:** 10.1101/2020.11.11.378117

**Authors:** Elina Zmeykina, Zsolt Turi, Andrea Antal, Walter Paulus

## Abstract

Sensorimotor mu-alpha rhythm reflects the state of cortical excitability. Repetitive transcranial magnetic stimulation (rTMS) can modulate neural synchrony by inducing periodic electric fields (E-fields) in the cortical networks. We hypothesized that the increased synchronization of mu-alpha rhythm would inhibit the corticospinal excitability reflected by decreased motor evoked potentials (MEP). In seventeen healthy participants, we applied rhythmic, arrhythmic, and sham rTMS over the left M1. The stimulation intensity was individually adapted to 35 ^mV^/_mm_ using prospective E-field estimation. This intensity corresponded to ca. 40% of the resting motor threshold. We found that rhythmic rTMS increased the synchronization of mu-alpha rhythm, increased mu-alpha/beta power, and reduced MEPs. On the other hand, arrhythmic rTMS did not change the ongoing mu-alpha synchronization or MEPs, though it increased the alpha/beta power. We concluded that low intensity, rhythmic rTMS can synchronize mu-alpha rhythm and modulate the corticospinal excitability in M1.

**Highlights:**

- We studied the effect of rhythmic rTMS induced E-field at 35 ^mV^/_mm_ in the M1
- Prospective electric field modeling guided the individualized rTMS intensities
- Rhyhtmic rTMS entrained mu-alpha rhythm and modulated mu-alpha/beta power
- Arrhythmic rTMS did not synchronize ongoing activity though increased mu-alpha/beta power.
- Rhythmic but not arrhythmic or sham rTMS inhibited the cortical excitability in M1

## 1. Introduction

Synchronously fluctuating transmembrane currents generate rhythmic activity in neural ensembles. These current flow alterations go along with the rhythmic shift between higher and lower excitability states [1].

Sensorimotor mu-alpha power in the range of 8-13 Hz reflects the corticospinal excitability fluctuation in the motor cortex. In humans, mu-alpha excitability can be measured non-invasively with the electro-/magnetoencephalogram (EEG/MEG) [2], or by recording motor evoked potentials (MEP) after transcranial magnetic stimulation (TMS). The amplitude of the MEP depends on the timing of the TMS pulse relative to the phase of the sensorimotor mu-alpha wave [3]. When TMS is delivered at the trough of the mu-alpha wave, the MEP amplitudes are larger than with TMS delivered randomly or at the peak [3].

Increased synchronization of the mu-alpha rhythm over central and parietal areas occurs during a motor inhibition task [4]. MEP amplitudes were reduced during the inhibition condition when participants had to inhibit the motor response as compared to the motor activation task and baseline [4]. These findings support the inhibition-timing hypothesis that assumes that mu-alpha oscillation is induced by inhibitory cells and reflects the shifts between the phases of maximal and minimal inhibition states [5].

Ten Hz rTMS tends to increase the corticospinal excitability in most participants [6]. This effect may appear opposite to the inhibition-timing hypothesis by which induced mu-alpha rhythm should inhibit rather than increase the corticospinal excitability. However, not one single parameter defines the neural mechanism of rTMS but rather the combination of the frequency and amplitude of stimulation and duration of the protocols.

We propose that one crucial parameter in deciding the effects of rTMS on corticospinal excitability level is the degree of mu-alpha synchronization. We predict that if 10 Hz rTMS can increase the degree of mu-alpha synchronization in the motor cortex, it should shift the oscillatory state into inhibition and temporarily decrease the corticospinal excitability. On the other hand, a 10 Hz protocol might perturb, rather than increase the degree of mu-alpha synchronization. This might be the case at high stimulation intensities or when using arrhythmic rTMS. By reducing mu-alpha synchronization cortical inhibition would be less and thereby it would result in a temporarily increase of corticospinal excitability.

In our previous study [7] we used a novel stimulation intensity selection approach for rTMS, which was based on prospective electric field (E-field) estimation. We observed ongoing parietal-occipital alpha synchronization at comparably low intensities in the range of 30-42% of the resting motor threshold corresponding to 35 and 50 ^mV^/_mm_ [7].

In the present study we extended our previous study by focusing on effects and aftereffects on sensorimotor mu-alpha synchronization and corticospinal excitability level. To this aim we applied rhythmic, arrhythmic, and sham rTMS protocols over the left primary motor cortex (M1) at 35 ^mV^/_mm_ E-field strength. We hypothesized that desynchronized mu-alpha activity would reflect a state of comparatively high excitability, whereas synchronized mu-alpha activity would reflect a state of inhibition and low excitability. We predicted that rhythmic rTMS would increase local mu-alpha synchronization and inhibit corticospinal excitability level, and hence reduce MEP amplitudes. On the other hand, we expected that arrhythmic rTMS would perturb mu-alpha oscillations and thereby lead to motor cortex excitation, i.e. to, increased MEP amplitudes.

## 2. Methods

### 2.1. Participants

Seventeen neurologically healthy volunteers (eight females) participated in the study. The age range was 24-32 years (mean ± SD: 27.4 ± 2.8 years). Although the dataset of one participant was incomplete we included the data in the analysis. Two participants did not complete the experiments due to a high resting motor threshold and non-tolerability of TMS. The sample size was determined based on earlier rTMS-EEG studies [8–10].

Before participation all volunteers filled out self-completed questionnaires to assess the study exclusion criteria. In cases of possible contraindications, a neurologist at the Department of Clinical Neurophysiology, University Medical Center Göttingen examined the volunteer. Inclusion criteria were no history or presence of medical, neurological, or psychiatric illnesses including epilepsy, drug and/or alcohol abuse, and no metal implants in the head, neck, or chest. We used the Edinburgh Handedness Inventory questionnaire to estimate the laterality index of participants. We included only right-handed participants with an index range of 30-100 (mean laterality index ± SD: 75 ± 15.5).

All participants gave written informed consent before participation. We performed all experiments according to relevant regulations. The Ethics Committee of the University Medical Center Göttingen approved the study (Application number: 36/4/19).

### 2.2. Overview of the experimental sessions

The design of the study is shown in Figure 1. In the first session we collected neuroimaging data to determine the stimulation target and prepare anatomically realistic head models for electric field simulations. In the second session we estimated the motor cortical target and the resting motor threshold. In the remaining sessions, we performed three courses of rTMS-EEG (i.e., rhythmic, arrhythmic, and sham protocols) in a randomized order. In each rTMS-EEG session, we assessed the immediate effects and the aftereffects of rTMS with EEG and MEP.

**Figure 1.**
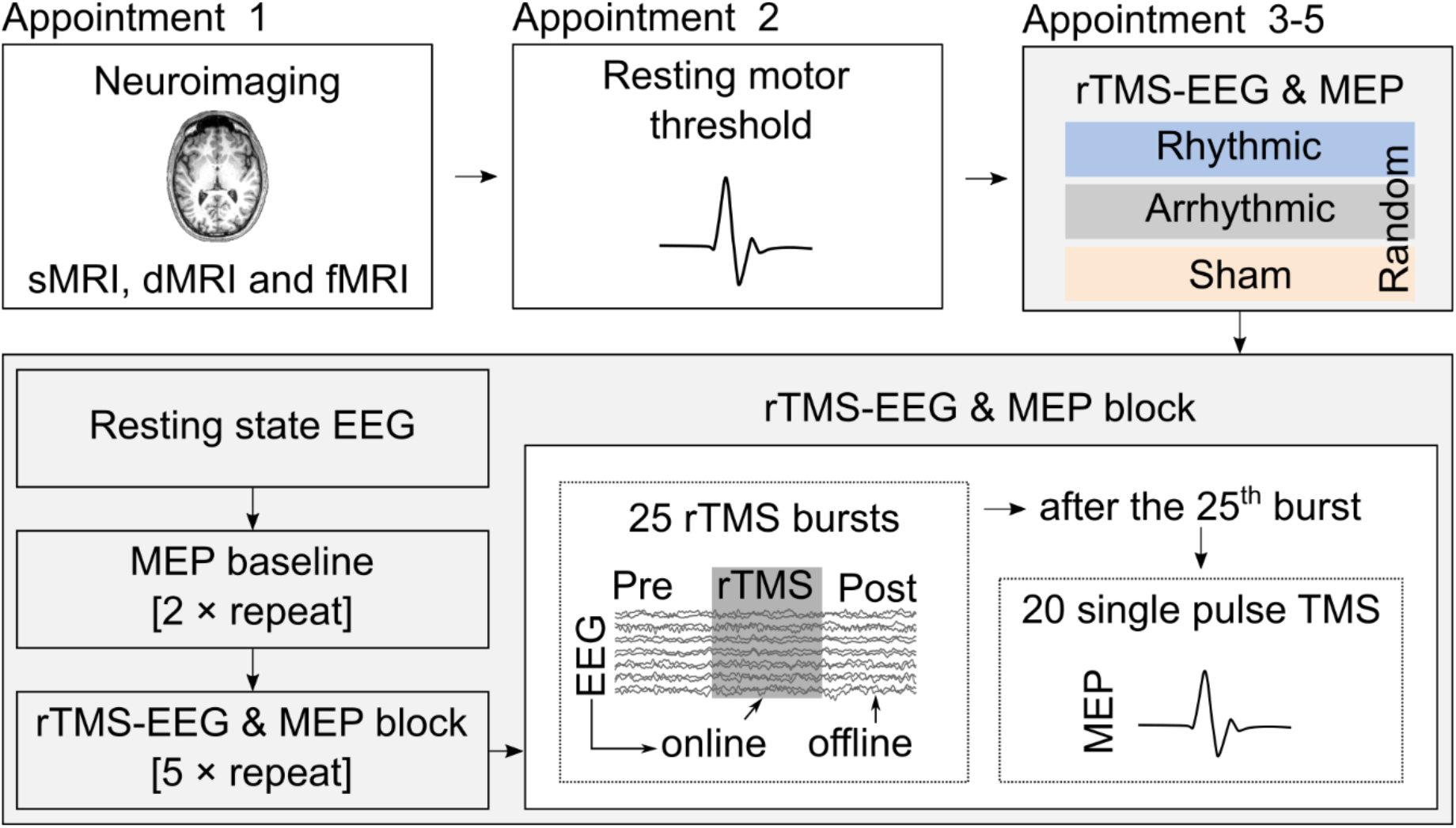
Schematic of study flow. After the neuroimaging and resting motor threshold estimation sessions, the participants took part in three rTMS sessions. Each session started with resting state EEG acquisition followed by determining the MEP baseline. Then, the participants received five rTMS-EEG and MEP blocks. In each block, we assessed the online and offline EEG effects. At the end of each block, we assessed cortical excitability by single pulse TMS.

### 2.3. TMS and neuronavigation

In both single-pulse TMS (spTMS) and rTMS the biphasic pulses were delivered using a MagPro X100 stimulator (MagVenture, Denmark) with a standard figure-eight coil (MC-B70), normal coil current direction, and 280 µs pulse duration. During stimulation, the participants sat in a comfortable chair with a chin and head fixator to minimize head movements. The TMS coil was placed over the motor cortex representation of the right first dorsal interosseous muscle, which we had previously identified as the highest local activation in the parametric t-map from the fMRI experiment.

To monitor the coil position during stimulation we used an MRI-based real-time neuronavigation system (Brainsight TMS Navigation, Rogue Resolutions Ltd) and coupled it with a Polaris Vicra infrared camera (NDI, Waterloo, Canada).

### 2.4. rTMS protocols

We performed three rTMS-EEG sessions with the individually predefined parameters of stimulation for the rhythmic, arrhythmic, sham stimulation protocols. The order of the protocols was randomized for each participant. The sessions were performed on three different days with at least 72 hours between each.

The rTMS bursts were delivered at the prospectively estimated individualized intensities (see Head modeling and E-field calculations). We set the intra-burst frequency of the rTMS bursts according to the individual alpha frequency (IAF) in the rhythmic and sham sessions. IAF was determined from the resting state continuous EEG data (see rTMS-EEG data acquisition). The intra-burst frequency used in the arrhythmic protocol was predefined by pseudorandomization of frequencies excluding 8-12 Hz and their harmonics [7].

In the rhythmic and arrhythmic protocols the coil position and angle were adjusted to the optimal for inducing the MEP response. In the sham protocol the coil was positioned at the M1 but was tilted by 90 degrees away from the head surface. In each protocol, a burst consisted of 20 TMS pulses with an inter-burst interval of 10 or 11 s. Each burst was repeated 25 times in a block. Each session consisted of five rTMS blocks, which resulted in 2,500 (= 20 × 25 × 5) pulses per session. For the stimulation parameters, see Table 1.

**Table 1.** Overview of rTMS stimulation conditions and stimulation parameters. Abbreviations: IAF – individual alpha frequency, MSO – maximum stimulator output.

|  | **Rhythmic** | **Arrhythmic** | **Sham** |
| --- | --- | --- | --- |
| Coil handle orientation | Parallel | Parallel | Perpendicular |
| Stimulation intensity | Individualized | Individualized | Individualized |
| Range of MSO % | 15 to 18 | 15 to 18 | 15 to 18 |
| Stimulation frequency | IAF | Pseudorandom | IAF |
| IAF Hz (mean ± SD) | 10.3 ± 0.9 | 10.2 ± 0.9 | 10.5 ± 0.6 |
| Pulses / burst | 20 | 20 | 20 |
| Number of bursts/block | 25 | 25 | 25 |
| Number of blocks/session | 5 | 5 | 5 |
| Total pulse number | 2,500 | 2,500 | 2,500 |

### 2.5. Head modeling and E-field calculations

We performed individual, anatomically realistic head modeling, and E-field calculations using the Simulation of Non-invasive Brain Stimulation (SimNIBS) software package [11]. We created the multi-compartment head models using the ‘mri2mesh()’ SimNIBS function based on T1-, T2-weighted images with and without fat suppression (see MRI and fMRI data acquisition). We assigned the standard conductivity values to the compartments [12], in S/m: scalp (0.465), bone (0.01), cerebrospinal fluid (1.654), gray matter (0.275) and white matter (0.126).

For each participant, we simulated the E-field induced at the target determined from the fMRI activation map with a standard figure-eight coil (MC-B70). The coil was positioned at 8.5 mm from the head surface (including the 8 mm EEG electrode thickness) with a handle direction of ca. 45° along the medium plane. We ran the calculations for intensities in the range of 15-20% of maximum stimulator output (MSO) and chose the intensity that induced around 35 ^mV^/_mm_ absolute peak E-field (99.9% percentile over the entire gray matter). The average intensity used for the rTMS protocols was 16.6 ± 1.1% MSO.

The simulation of E-field was performed twice: before and after rTMS-EEG sessions. The second estimations of E-field were performed after stimulation using session specific coil position information exported from the Brainsight TMS Navigation software. The MNI coordinates of coil location and direction were transformed into subject-specific space using the ‘mni2subject_coords()’ SimNIBS function.

### 2.6. Data acquisition

#### 2.6.1. MRI and fMRI data acquisition

We acquired anatomical, diffusion-weighted, and functional magnetic resonance imaging data with a 3T MRI-scanner (Siemens Magnetom TIM Trio, Siemens Healthcare, Erlangen, Germany). Functional MRI was collected during rhythmic, stereotypic movement using the first dorsal interosseous muscle, i.e. movement of forefinger from side to side. The fMRI data preprocessing was performed with the Statistical Parametric Mapping (SPM 12, Welcome Department of Imaging Neuroscience, London, UK) software package implemented in MATLAB software. Following preprocessing described elsewhere [7], the general linear model was applied at the single-subject level. Voxels were identified as significant if p < 0.05 (family-wise error corrected for multiple comparisons on the voxel level). The individual activation T-map as well as T1 image were uploaded to the neuronavigation system.

#### 2.6.2. EMG acquisition

We recorded the EMG with an Ag–AgCl electrode pair attached to the first dorsal interosseous muscle of the right hand in a belly–tendon montage. Signals were sampled at 5 kHz, amplified and bandpass filtered between 2 Hz and 4 kHz, and digitized using a 1401 AD converter (CED 1401, Cambridge, UK). All EMG measures were recorded with Signal software (CED, version 4.08).

The experiment started by determining the resting motor threshold (RMT) at the hotspot. The initial position was defined as that with the fMRI local activation maximum derived from the parametric t-map at the anatomical hand knob formation. The spTMS was delivered at 0.25 Hz by placing the stimulation coil orthogonally to the central sulcus at 30% MSO. We increased the intensity in increments of 2% MSO until the stimulation evoked MEPs. We further decreased or increased intensity by 1% MSO until we identified the lowest intensity that evoked at least five out of ten MEPs with a peak-to-peak amplitude > 50 µV. The average RMT was 44.0 ± 8.5% of MSO.

In the rTMS-EEG sessions we determined the 1 mV MEP threshold that we used for the baseline MEP measurements and for assessing the corticospinal excitability level after rTMS. To find the 1 mV threshold, we delivered 20 TMS pulses initially at 120% of RMT. We then averaged the peak-to-peak MEP amplitudes and evaluated whether the 1 mV threshold was reached. In the next block, we decreased or increased the TMS intensity by 5% RMT, if necessary. We repeated this procedure until we obtained a peak-to-peak MEP amplitude within the range of 0.8-1.2 µV.

We performed the baseline measurements twice to ensure that the baseline MEP amplitudes were accurately assessed. In each measurement, we delivered 20 TMS pulses. Following the baseline measurement and immediately after each rTMS block, we assessed the MEPs using the 1 mV threshold intensity.

#### 2.6.3. rTMS-EEG data acquisition

##### EEG acquisition

In each rTMS-EEG session we recorded the EEG simultaneously with rTMS from 64 Ag/AgCl active EEG electrodes (actiCAP slim, BrainVision LLC, Germany) at a 2.5 kHz sampling rate without hardware filters (actiChamp, Brain Vision LLC, Germany). Ground and reference electrodes were located at AFz and FCz, respectively. Impedance values were maintained below 20 kΩ.

Stimulation sessions started with recording two blocks of continuous resting state EEG for four minutes with eyes open and eyes closed. During the recording we instructed the participants to sit calmly and relaxed, minimize blinking and horizontal eye movements, not to move their arms, legs, or facial muscles, to avoid any calculations or repetitive mental activity such as reproducing any texts, lyrics, or melodies. The participants wore QuietControl 30 wireless headphones with active noise reduction and white noise masking during all recordings (Bose Corporation, USA). The volume level was always kept below the manufacturer’s recommended safety limits. This procedure minimized but did not eliminate the sound produced by the TMS stimulus. Therefore, at the end of each session participants were asked to evaluate the click sound by scale from -100 to +100 percent, where zero meant the same loudness of click and white noise. The coordinates of each electrode were saved to the neuronavigation system once at the end of the final rTMS-EEG session.

We performed offline data analysis with the FieldTrip toolbox for EEG-and MEG analysis [version 20170119; 85; http://fieldtrip.fcdonders.nl] as described in reference [7]. We determined the global peak alpha frequency in the range of 8-12 Hz in the eyes open state. For three participants the peak frequency during the eyes open state was not clearly defined. We, therefore, chose the stimulation frequency from the global peak alpha defined from the eyes-closed state.

The EEG data of each participant consisted of resting-state EEG at the beginning and the end of each session; 140 trials of spTMS (40 trials from baseline measurements and 20 trials after each rTMS block) and 125 trials of rTMS-EEG recording.

### 2.7. Data analysis

#### 2.7.1. E-field analysis

Descriptive statistics of the absolute E-fields and its normal component were performed in MATLAB for three predefined anatomical regions of interests (ROIs), i.e., the left precentral and left postcentral gyri, and left central sulcus. From each ROI, we extracted the robust minimum (0.1%) and maximum (99.9%) values, as well as the global mean.

#### 2.7.2. MEP analysis

EMG recordings were converted with cfs2mat utility (https://github.com/giantsquidaxon/cfs2mat) to MATLAB ‘.mat’ format where they were further preprocessed. The data were visually inspected for activation higher than 50 µV in the 80 ms pre-TMS interval. Trials containing pre-activation were excluded from further analysis. Altogether 2.1 % of all MEP trials were excluded from the dataset.

Peak-to-peak MEP amplitude was calculated for 10-50 ms after the TMS pulse. We transformed the MEP data using Matlab’s base 10 logarithm function ‘log()’ to reduce the frequency and the weight of outliers, and to improve variance homogeneity across experimental conditions. After log transformation we removed four additional data points because they exceeded ± 3 SD of the global mean value.

The MEP amplitudes were averaged across the blocks at the time points. The effect of rTMS protocols on cortical excitability was tested using the linear mixed-effect model implemented in R (version 4.3.0), R-Studio integrated development environment (version 1.3.1093), and ‘lmerTest’ package (version 3.1.2) [14–16].

The base model only included the random intercept for the participant using the formula described in Table 2. In the next model we added the effect of Protocol (three levels: rhythmic, arrhythmic, and sham rTMS) and Time (six levels: baseline and five subsequent measurements).

**Table 2.**
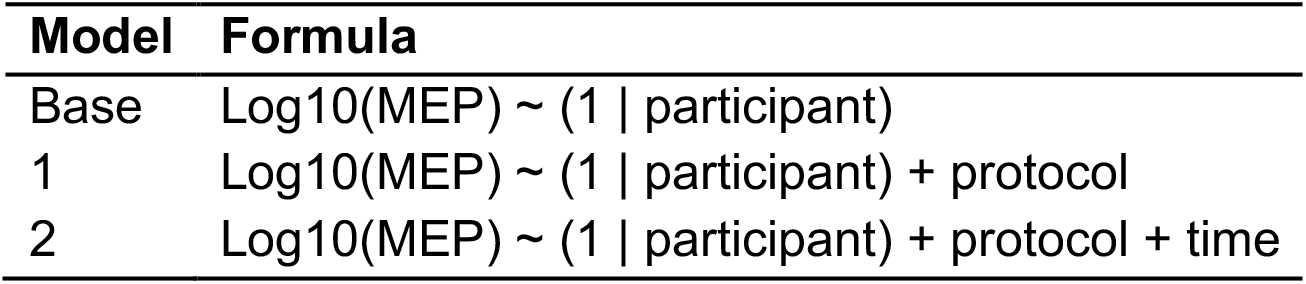
The formulae of the tested models.

We used the ‘anova()’ function to test which model provided the best parsimonious data fit. Models with a p-value of 0.05 or less were considered to be significantly better than the previous model. On the winning model, we ran the ‘anova()’ function to perform Type III Analysis of Variance with Satterthwaite’s method.

#### 2.7.3. RTMS-EEG analysis

##### EEG preprocessing

**The** EEG data were preprocessed offline in MATLAB (2017b, Mathworks) using Fieldtrip toolbox (v.20180114, [13]) and custom-written code. The data were cleared of TMS-induced artifacts by the same procedure as described in [7]. The data were then re-referenced to a common reference and were cut into segments of 8.5 sec length. Trials were aligned to the last TMS pulse of the burst and contained 5.5 sec before and 3 sec after the pulse.

The data were visually inspected to identify and remove excessively noisy channels and trials with jumps or muscle artifacts. The data were resampled to 1250 Hz, and the signals relating to eye blinks and eye movements were identified and removed by the second ICA. On average, 15.8 ± 4 (R); 16.7 ± 3.6 (AR); 15.8 ± 4.2 (SH) components and 6.2 ± 2 (R); 4.2 ± 1.3 (AR); 3.8 ± 1.5 (SH) channels were removed from the data. The remaining clean dataset contained on average 103 ± 6.5 (R); 107.8 ± 9 (AR); 101.7 ± 13.5 (SH) trials.

##### Phase-locking value (PLV)

We calculated the degree of synchronization between the ongoing signal phase and the TMS pulses by estimating the PLV during the rTMS burst [17]. We used our previous analytical pipeline described elsewhere [7]. Briefly, we simulated the sinusoidal wave at the stimulation frequency and aligned its phase to TMS pulses. The simulated signal was appended with cleaned EEG data as an additional channel. The data was transformed by complex Morlet wavelet decomposition from 1 to 25 Hz.

PLV was computed between the phases of the original signal and the simulated wave. The PLVs were normalized relative to the baseline at 500 ms before TMS bursts onset. Then we focused the analysis on the online effect which showed the changes in PLVs during stimulation and the offline effect which showed PLV changes immediately after TMS bursts. We averaged normalized PLVs separately for online and offline effects over time from -2 to 0 sec and from 0 to 2 sec correspondingly. They were further compared between protocols by group-level, nonparametric, cluster-based permutation test (10,000 permutations, two-tailed, significance accepted at p < 0.05). We performed two dependent t-tests to compare rhythmic with arrhythmic and sham separately for online and offline effects.

##### Power analysis

Time-frequency transformed data were further investigated for power changes. Residual TMS artifacts could affect the signal amplitude and power [7], and we, therefore, performed power analysis on artifact-free intervals from 0.2 to 2 sec after an rTMS burst. The power was extracted as a real part of complex Wavelet decomposition and averaged over all channels for the three protocols. The trials were split for ‘before’ and ‘after’ intervals that corresponded to 1.8 sec prior and post rTMS. The difference in power was statistically tested using activation versus baseline t-statistic (‘actvsblT’ in Fieldtrip). The analysis was performed on all sensors for 5-25 Hz frequency. Statistical significance was assessed by nonparametric, cluster-based permutation test (10,000 permutations, two-tailed, significance accepted at p < 0.05).

Next, we investigated the single-trial alpha power by reconstructing the virtual source signal at the cortical level. For that we created a subject-specific virtual channel using the linear constrained minimum variance (LCMV) beamformer approach [18]. The coordinates of the source corresponded to the ‘hotspot’ of the stimulation site, which was exported from E-field estimations (see section 2.5.1) in subject-specific space.

A realistic three-layer volume conduction model was constructed using the individual MRI using the boundary element method [19]. A grid with 10 mm^2^ resolution was created per individual, which was subsequently normalized to MNI space. The spatial filter was constructed from the full trial length and then was used to reconstruct the virtual source signal. Time-frequency decomposition was estimated for 1-30 Hz using complex Morlet wavelet. The wavelets contained seven cycles with a three-Gaussian window. The decomposition was performed for the trial interval from 3.5 before to 2.5 sec after the TMS offset. The virtual source power was compared between active protocols (rhythmic, arrhythmic) versus sham by cluster-based permutation statistical test (10,000 permutations, two-tailed, significance accepted at p<0.05).

## 3. Results

In the present study, we assessed the effects on the EEG during rTMS bursts and the aftereffects in the inter-burst intervals, i.e., immediately after the rTMS bursts. We applied phase and power-based analysis and reconstructed the signal of the virtual source placed in the grey matter of the stimulated target. Moreover, we tested the aftereffects of rTMS on the corticospinal excitability level by analyzing the size of the MEP amplitudes after the end of the rTMS block, i.e., after 25 rTMS bursts.

### 3.1. Increased mu–alpha synchronization during and after rhythmic rTMS

First, we characterized the amount of synchronization during the rTMS bursts (online) and immediately after the stimulation cessation (offline; see Figure 2). We estimated PLVs that represent the amount of consistency of the EEG signal phase-locked to the external TMS pulses [20].

During the rTMS bursts (online), a nonparametric cluster-based permutation statistical test revealed a significant increase in PLVs in the mu-alpha range 8-13 Hz comparing the rhythmic versus arrhythmic (p < 0.001) and rhythmic versus sham (p < 0.001) rTMS protocols (see Figure 2A). In the rhythmic versus arrhythmic comparison, we found the largest difference in PLV increase in the posterior (T_peak_(14) = 4.27) and the left frontal (T_peak_ (14) = 4.46) regions (see Figure 2B, left). When rhythmic rTMS was compared to sham rTMS protocol, we found the highest PLV increase in the left central region (T_peak_ (13) = 4.03; see Figure 2B, left). Rhythmic rTMS synchronized ongoing sensorimotor mu-alpha rhythms as indicated by increased phase-locking values.

**Figure 2.**
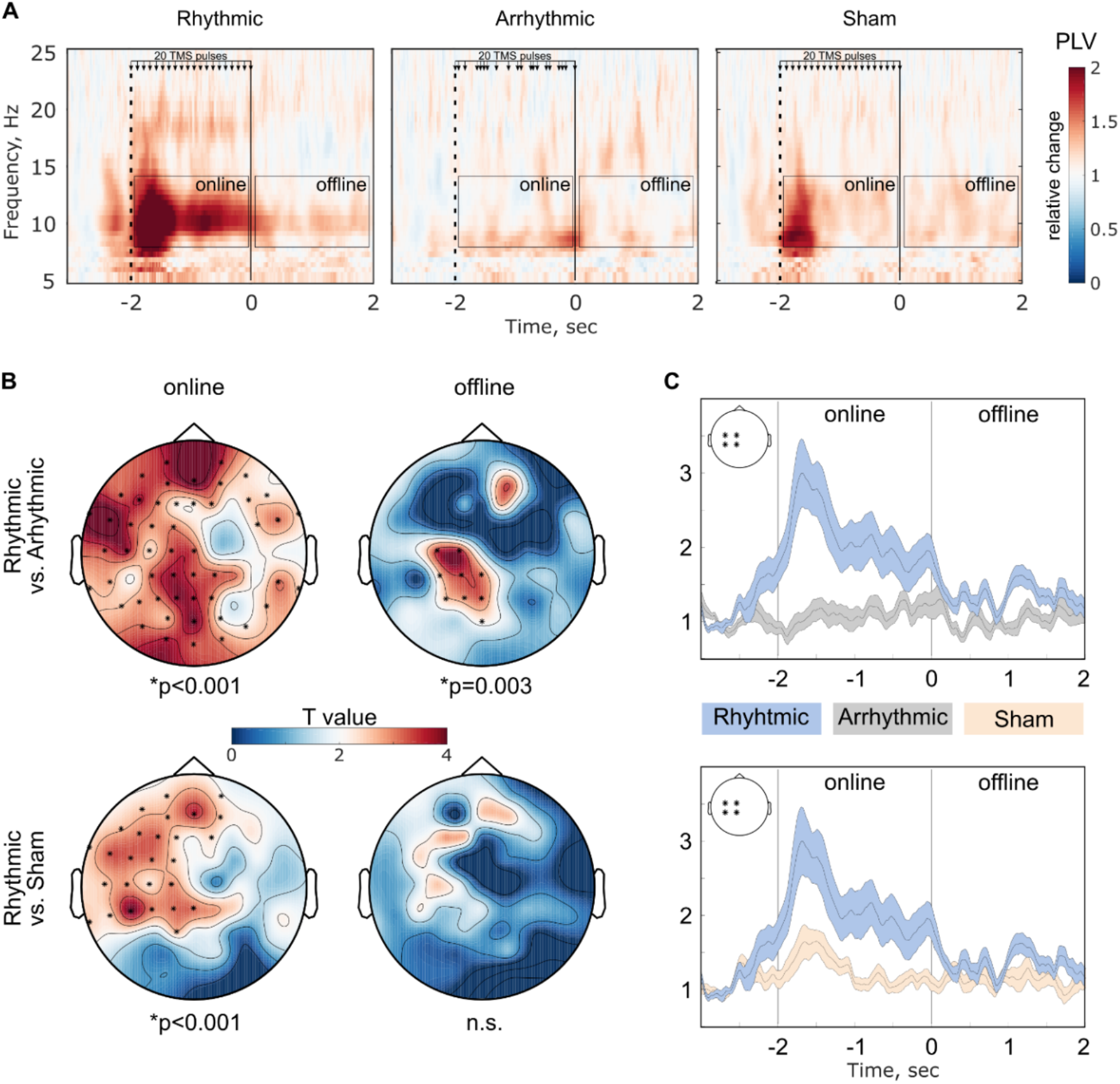
Rhythmic rTMS synchronized ongoing mu-alpha rhythms indicated by increased phase-locking values near the stimulation target. A) Global time-frequency representation of PLVs during and after rTMS burst for rhythmic, arrhythmic, and sham protocols (from left to right). PLVs are normalized by 500 ms interval before rTMS onset. B) Statistical tests revealed a significant increase of mu-alpha synchronization during rhythmic rTMS compared with arrhythmic or sham protocols. Immediately after rhythmic rTMS burst, mu-alpha synchronization is increased compared wiht arrhythmic but not with sham protocol. C) Time course of PLVs averaged over four left central electrodes (C1, C3, CP1, CP3).

Immediately after rTMS bursts (offline) we found a significant difference only between the rhythmic and arrhythmic protocols (p = 0.003) with a peak t-value over the central electrodes (T_peak_ (13) = 3.71) on the stimulation site (see Figure 2B, right and Figure 2C). The difference between rhythmic versus sham was not significant.

### 3.2. Rhythmic rTMS increased alpha/beta power after stimulation

We then investigated mu-alpha power changes induced by rTMS protocols. We estimated the relative change of global power by time-frequency transformation. The mu-alpha power was statistically compared between ‘after’ and ‘before’ EEG intervals. These intervals were free of any residual TMS artifacts. Statistical tests revealed significant clusters for rhythmic (T_peak_ (14) = 1.64, p = 0.01), arrhythmic (T_peak_ (13) = 1.87, p = 0.002) condition but not for the sham (T_peak_ (14) = 0.99, p = 0.24) protocol (see Figure 3).

**Figure 3.**
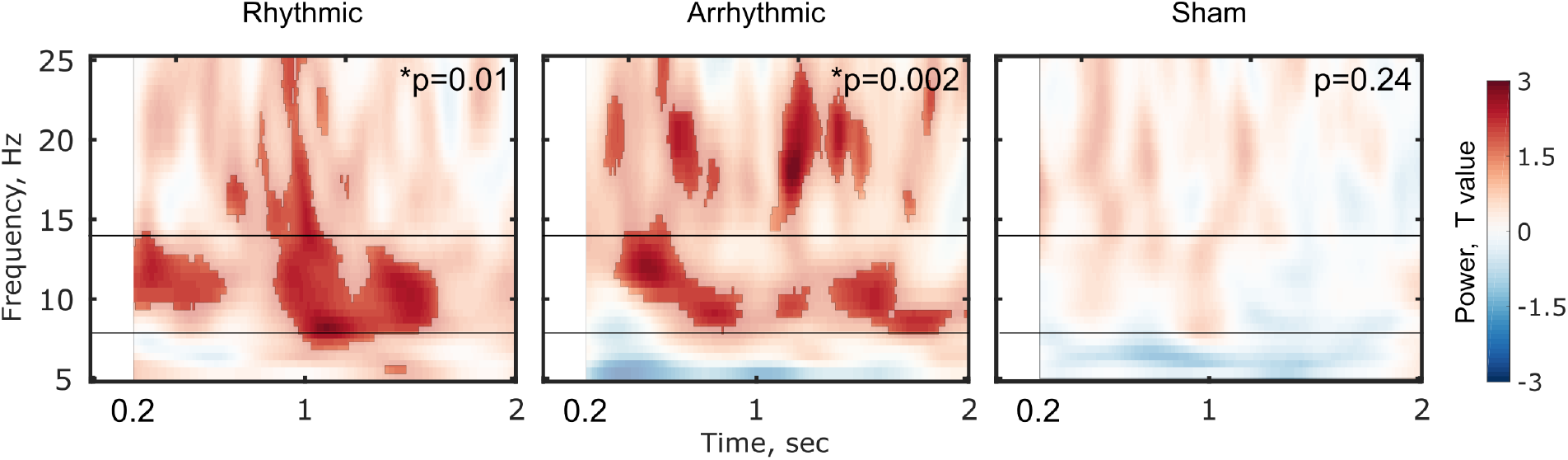
Active stimulation protocols (rhythmic and arrhythmic) increased power in mu-alpha and beta frequency ranges immediately after stimulation (inter-burst intervals). Sham rTMS applied at IAF did not change the power of mu-alpha rhythm. Time-frequency plots are masked with p<0.05.

To investigate the location-specific power change we projected the sensor level EEG to the source space by reconstructing the virtual channel signal (see Figure 4). The source location was selected on the cortex surface with coordinates corresponded to the peak E-field, which were estimated individually (see Figure 4B). The statistical test of time-frequency transformation of the virtual channel revealed a significant increase in the mu-alpha (T(14) = 0.93, p = 0.040) and beta (T(14) = 1.05, p = 0.032) power (see Figure 4C). Although the arrhythmic protocol increased the global and virtual source power, it was not significantly different from the sham protocol (T(13) = 0.75, p = 0.065; see Figure 4C).

**Figure 4.**
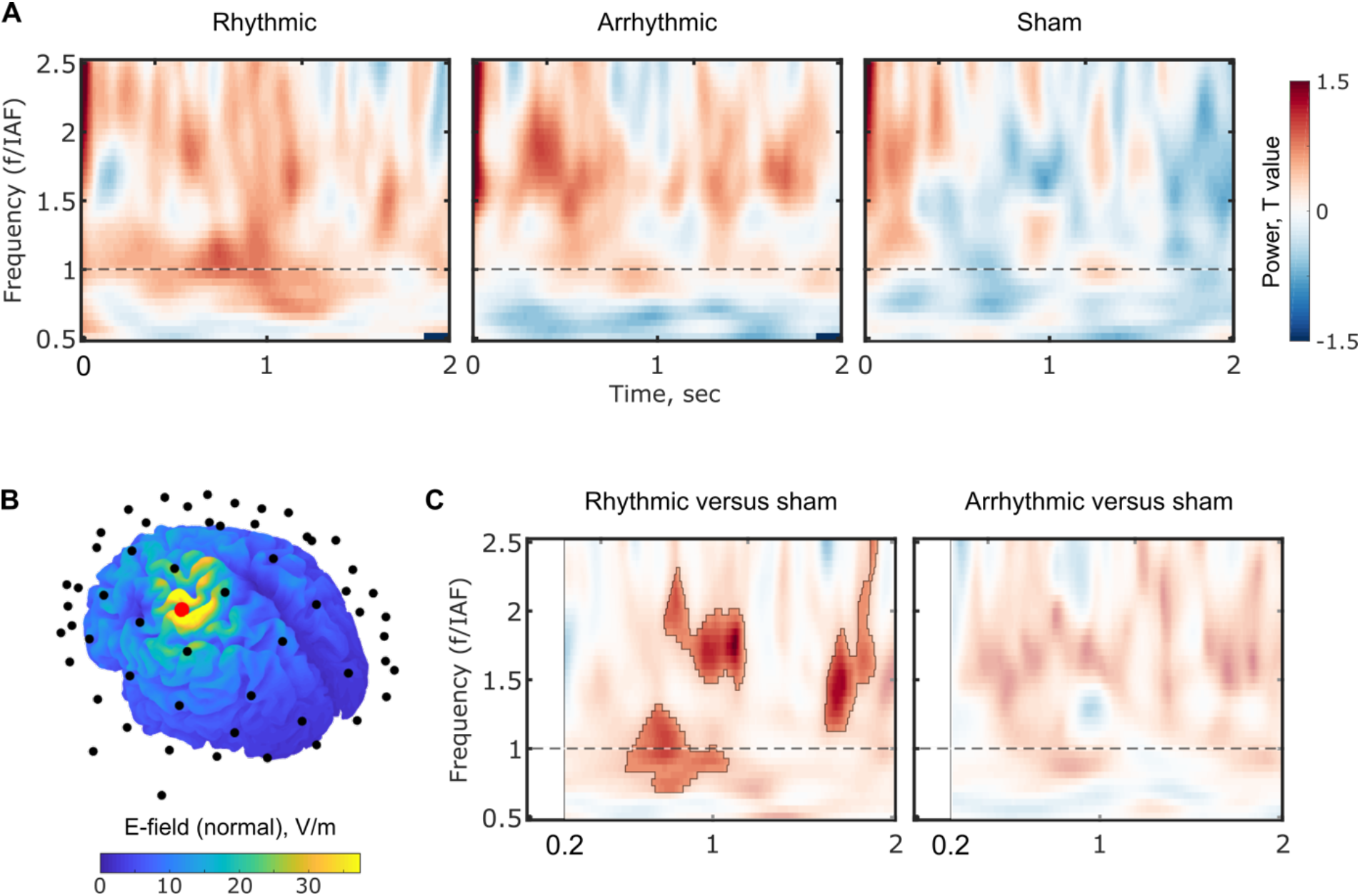
Rhythmic rTMS protocol increased local mu-alpha and beta power at the stimulation target. A) Time-frequency representation of relative power. The frequency scale is normalized to stimulation frequency (at IAF). Value one on the ordinate corresponds to IAF. B) Example of source localization for a single participant. The head model and E-field estimations were used for locating the virtual source in the peak E-field induced by TMS shown as a red sphere. Black dots show the location of the EEG electrodes. C) Statistical maps of the time-frequency representation for the virtual source. Significant clusters (p < 0.05) are marked by contour lines.

### 3.3. Rhythmic rTMS decreased corticospinal excitability level

We assessed the changes in cortical excitability by focusing on the log-transformed peak-to-peak MEP amplitudes. The winning model included the random intercept for participants and the fixed effect for the Protocol (*χ*^2^ 1, *N* = 3 = 9.9466, p = 0.0069), the latter of which had a significant main effect (*F*(2, 244.33) = 5.0113, *p* = 0.0072). Further analysis revealed that relative to sham stimulation, rhythmic rTMS significantly reduced the MEP amplitudes (*t* = −3.171, *df* = 243.27, *p* = 0.0017). On the other hand, arrhythmic rTMS had no significant effect on the MEP amplitudes (*t* = −1.467, *df* = 244.85, *p* = 0.1435). The Bonferroni-adjusted alpha levels were 0.0034 and 0.2870 for the rhythmic and arrhythmic protocols, respectively.

Then, we focused on the rhythmic protocol and studied the correlations between the MEP amplitudes and the degree of mu-alpha synchronization and the spectral power changes (see Figure 5). None of the correlations were significant (all p-values > 0.355).

**Figure 5.**
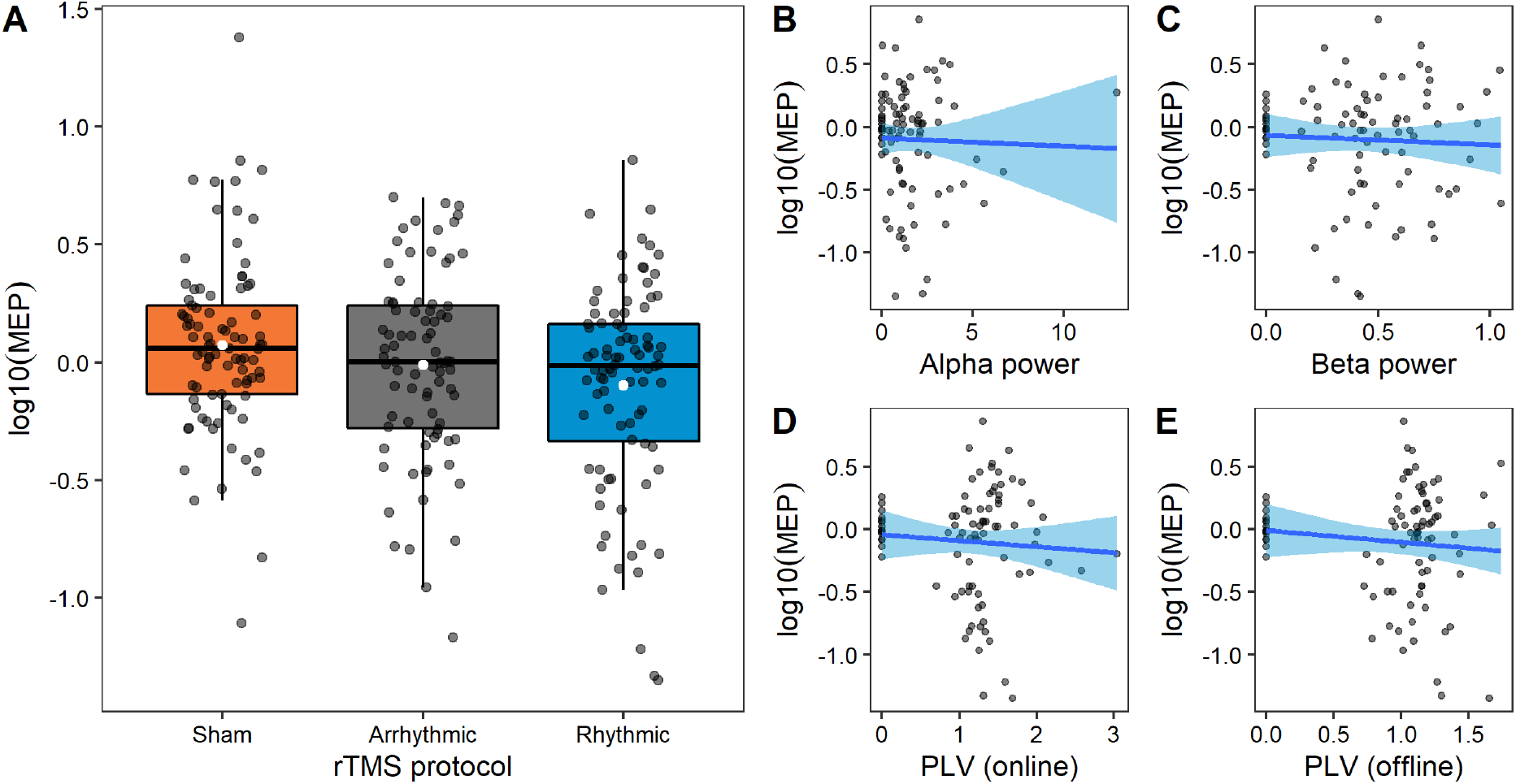
Rhythmic rTMS decreased the MEP amplitude. A) Log-transformed MEP amplitudes according to the rTMS protocols. White dot shows the mean values, black dots represent individual measurements. B-E) Linear relationships between the individual log-transformed MEPs and the individual EEG parameters. Alpha and beta power values (top) were extracted from virtual source time-frequency analysis as a peak power (normalized to baseline) for mu-alpha (IAF ± 1Hz) and beta rhythm (15-20 Hz) for 0.2-2 sec after rhythmic rTMS bursts. PLV values (bottom) are averaged over four channels (C1, C3, CP1, CP3) at the stimulation location at the IAF ± 1Hz.

## 4. Discussion

In the present study, we demonstrated that low-intensity rTMS over the left M1 influenced both oscillatory activity and corticospinal excitability level that outlasted the stimulation period. Furthermore, we replicated our previous findings on entrainment of parietal-occipital alpha rhythm by weak rTMS-induced E-fields [10], now at the motor cortex, and we extended our previous work by showing that rhythmic rTMS enhanced sensory-motor mu-alpha rhythm and decreased corticospinal excitability.

### 4.1. Electrophysiological effects of low-intensity rTMS

One question of key importance in applying rTMS is the selection of the stimulation intensity [15]. In the present study we used the prospective E-field estimation approach where we defined the stimulation intensity using computational models of the rTMS induced E-field. We showed that low-intensity rTMS at an individual mu-alpha frequency over the left occipital cortex [10] and now over the left motor cortex increased ongoing neural synchrony.

We extended the EEG analysis to the induced effects on the phase and power of the mu-alpha rhythm after the rTMS bursts. We found that the rTMS-increased neural synchrony in M1 lasted for up to two seconds after applying rhythmic but not arrhythmic or sham rTMS. The explored intervals were free from the TMS-produced artifacts, such as decay, ringing, or muscle artifacts.

Moreover, the degree of mu-alpha synchrony was significantly higher after rhythmic than after arrhythmic rTMS, though they were not significantly different compared with sham rhythmic rTMS. This finding most likely indicates that the sound from rhythmic TMS clicks contributed to the increase in mu-alpha synchronization [21]. However, the increase after sham was much lower than after real rTMS in both online and offline intervals. Therefore, we believe that the observed effect did not originate solely from the rhythmic rTMS clicks sound. Moreover, we did not find mu-alpha synchronization in the temporal electrodes, which would indicate the entrainment of mu-alpha rhythm through auditory input. Based on these grounds, we conclude that rhythmic rTMS can induce and maintain the synchronized oscillatory activity at the stimulation target.

Furthermore, we investigated global and local power changes for up to two seconds after the rTMS bursts. We observed increased global alpha power following rhythmic and arrhythmic but not sham rTMS. Here the maintained oscillatory activity resulted not only in the phase-locking changes but also in the increased power of mu-alpha (8-14 Hz) and beta (15-20 Hz) frequency ranges.

This was not the case for local alpha power at the source level where we found the alpha and beta power increase only in rhythmic rTMS. Our findings are in line with previous studies where participants received conventional stimulation intensities of rTMS (80-100% of resting motor threshold) applied over the left M1 at rest [22,23]. The authors found a significant difference between the real and sham rTMS in mu (10-12 Hz) and beta (13-30 Hz) power for 5 sec after real rTMS trains.

### 4.2. Arrhythmic rTMS induces alpha perturbation in the occipital but not the motor cortex

One interesting finding is that arrhythmic rTMS induced different aftereffects in the occipital and sensorimotor mu-alpha rhythms [24]. Whereas arrhythmic rTMS significantly suppressed parietal-occipital alpha [24], in the present study it increased the sensorymotor mu-alpha power (see Figure 3). Apart from the stimulation target we used closely matched stimulation parameters in the two experiments. On the other hand, rhythmic rTMS increased the alpha power in both experiments.

The reason for this finding is currently not well understood. One crucial difference between the studies was that the amplitude of alpha rhythm was stronger and the peak alpha frequency was more prominent at the occipital than at the sensorymotor area. We speculate that the properties of the endogenous oscillation could have shaped the direction of the electrophysiological response to arrhythmic rTMS. In cortical regions with pronounced peak alpha frequency, arrhythmic perturbation suppressed alpha power. On the contrary, in cortical regions with less pronounced peak alpha frequency, arrhythmic rTMS increased alpha power. Nevertheless, further studies are needed to better understand the neural mechanisms of arrhythmic rTMS on ongoing oscillatory activity in different cortical regions.

### 4.3. Low-intensity rTMS affects corticospinal excitability

We also demonstrated that rhythmic rTMS inhibited the corticospinal excitability level as indicated by the reduced peak-to-peak MEP amplitudes. Contrary to our hypothesis, arrhythmic rTMS did not result in an MEP amplitude increase; we found no significant difference from the sham condition.

The observation that only rhythmic but not arrhythmic rTMS induced aftereffects in corticospinal excitability argues for the role of spike-timing-dependent plasticity. This form of Hebbian plasticity emerges by synchronously activating pre- and postsynaptic neurons. Because arrhythmic rTMS cannot achieve this tight temporal correlation between pre- and postsynaptic neurons, it did not induce lasting changes in corticospinal excitability.

In addition, we explored the relationships between MEP amplitudes and the EEG parameters such as the phase-locking value during and after the rTMS bursts as well as peak mu-alpha and beta power at the stimulation target. However, we found that changes in MEP amplitudes after the rTMS block did not correlate with changes in the EEG during or shortly after the rTMS bursts.

Several rTMS studies have demonstrated increased MEP amplitudes after applying 10 Hz rTMS over the primary motor cortex [6,26–28], whereas others reported decreased MEP amplitudes or no aftereffect [6,10,22,27–29]. Moreover, MEP amplitudes and EEG readouts correlated only weakly (CorrCoef < 0.1) [30,31]. Therefore, it is likely that these outcome measures do not reflect the same neural mechanisms.

### 4.4. Conclusions

In summary, we found that rTMS applied at an intensity of ∼40% RMT is effective to induce changes in ongoing electrophysiological and aftereffects in corticospinal excitability. The stimulation intensity used in the current study and in our previous [10] studies (38.8 ± 6.5% RMT) was approximately half of that applied in conventional rTMS studies, i.e., 80-120% of the motor threshold [32]. However, more research on this topic needs to be undertaken before the association between electrophysiology and cortical excitability is more clearly understood.

## 5. Authors’ contribution

EZ: conceptualization, study design, project administration, investigation (literature search, data collection), software, formal analysis, visualization, data interpretation, data curation, and writing the original draft

ZT: conceptualization, methodology, formal analysis, visualization, data interpretation, and writing the original draft

AA: data interpretation, and writing the original draft

WP: conceptualization, study design, funding acquisition, resources, data interpretation, and writing the original draft

## 6. Acknowledgments

We would like to thank Prof. Thomas Crozier for his comments on the manuscript.

